# Effects of stochasticity on the length and behaviour of ecological transients

**DOI:** 10.1101/2021.03.25.437077

**Authors:** Alan Hastings, Karen C. Abbott, Kim Cuddington, Tessa Francis, Ying-Cheng Lai, Andrew Morozov, Sergei Petrovskii, Mary Lou Zeeman

## Abstract

There is a growing recognition that ecological systems can spend extended periods of time far away from an asymptotic state, and that ecological understanding will therefore require a deeper appreciation for how long ecological transients arise. Recent work has defined classes of deterministic mechanisms that can lead to long transients. Given the ubiquity of stochasticity in ecological systems, a similar systematic treatment of transients that includes the influence of stochasticity is important. Stochasticity can of course promote the appearance of transient dynamics by preventing systems from settling permanently near their asymptotic state, but stochasticity also interacts with deterministic features to create qualitatively new dynamics. As such, stochasticity may shorten, extend, or fundamentally change a system’s transient dynamics. Here, we describe a general framework that is developing for understanding the range of possible outcomes when random processes impact the dynamics of ecological systems over realistic time scales. We emphasize that we can understand the ways in which stochasticity can either extend or reduce the lifetime of transients by studying the interactions between the stochastic and deterministic processes present, and we summarize both the current state of knowledge and avenues for future advances.

## 1 Introduction

Two major goals of ecological theory are to make predictions, and to explain past observations. In both cases, qualitative changes in dynamics through time represent both a challenge and an opportunity. For prediction, a sudden change in dynamics is important to capture. In parallel, understanding the limits to prediction, in time or in other ways, is important. Both for prediction of the future and for understanding the processes that lead to the current state of the system, the presence of large changes in dynamics, sometimes referred to as black swan events [1, 2], presents a challenge. How can these events be understood using ecological models?

There is increasing recognition that transients can play a critical role in ecological systems [3, 4, 5, 6, 7, 8, 9, 10], building on the variety of long transient behaviors exhibited by nonlinear dynamical systems [11, 12]. For example, regime shifts are an important phenomenon in ecology, in which the system behavior changes suddenly without any warning (e.g., sudden species extinction) [13, 14, 15]. The traditional view is that regime shifts are caused by parameter drifting. However, as recently emphasized, even without any parameter change, transient behaviour can lead to regime shifts [8, 9].

Earlier work has emphasized the possibility of sudden changes in dynamics even in deterministic models with constant parameters [8], but stochasticity is ubiquitous in real ecosystems and will affect transients [10]. How has the deterministic view limited our understanding of sudden shifts in ecosystems, and how does this understanding deepen when we account for stochasticity in our theoretical constructs and models? This question, in the context of observations of changing ecological dynamics and black swan events in ecology [1, 2], fits in with the recent recognition of the importance of focusing on dynamics on ecological time scales. Stochasticity can play an important role in determining dynamics on realistic time scales.

Real-world ecosystems are subject to inevitable and constant influences of stochastic disturbances that can have significant effects on the population dynamics [16, 17, 18, 19, 20, 21, 22, 23, 24, 25, 26, 27, 28, 29, 30, 8]. A particularly notable example includes the population dynamics of Dungeness crab, *Cancer magister,* along the USA West Coast [31]. In this system, chaotic-like oscillations were analyzed using a method that combined data analysis and modelling fitted from data to reveal that the oscillations were actually long transient relaxations due to stochastic perturbations of a stable equilibrium. In addition, random perturbations of cyclic population dynamics can also result in a chaotic-like behaviour, which was observed in the experimental dynamics of *Tribolium* [32]. Although there have been many individual, well-studied examples illustrating these points, a recognition of common themes arising in a discussion of stochastic transients in ecological systems reveals both new insights into ecological dynamics and suggests important future research directions.

We start from the premise that in natural systems, noise and random disturbances are inevitable. We consider noise that affects one or more state variables, perturbing them with some magnitude, direction, and frequency. Noise can influence long transients in a variety of ways (figure 1). Noise may certainly alter long transient dynamics that were created by another mechanism and already present in the ecological system. Importantly, stochasticity can also provide an alternate mechanism for long transient dynamics, creating a long transient that would not otherwise occur. There are two major types of stochasticity in ecological systems: external perturbations due to random variations in the environmental conditions, and internal population fluctuations. Some environmental stochasticity can be modeled as additive Gaussian white noise [33, 34], while internal stochasticity is effectively demographic noise [35, 23, 36, 37] that needs to be described as multiplicative noise with its strength depending on the fluctuating abundance variable. Demographic noises are thus correlated, colored stochastic processes.

**Figure 1:**
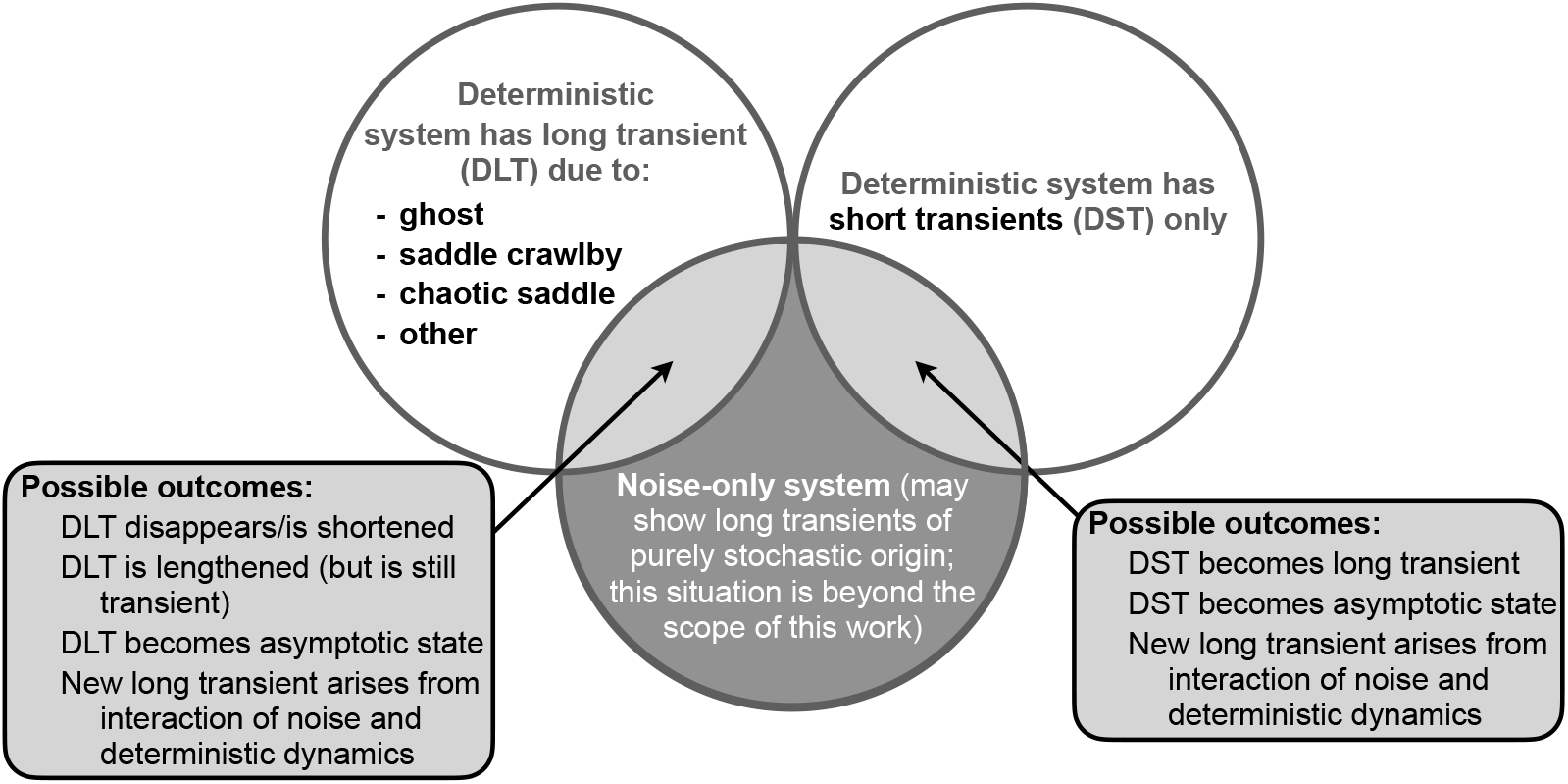
Real world dynamics fall into the light gray regions, where deterministic and stochastic processes interact. The length and nature of transient dynamics in these regions depends both on the presence of deterministic features known to promote long transients and on properties of the noise. DLT = deterministic long transient (i.e. a long transient that exists in the deterministic part of the dynamics); DST = deterministic short transient. The definition of a ‘long’ transient can be found in Section 3, and for simplicity we refer to all other transients as ‘short’.

We begin this exploration of the role of noise in creating and influencing long transients with some simple examples, illustrating the important point that stochasticity can either extend or reduce transients. Using the simple examples as a jumping off point, we then undertake a systematic exploration of transients in nonlinear (density-dependent) ecological systems. Even these simple examples bring out the important point that the definition of a transient for a stochastic system may be less clear cut than for a deterministic one. In particular, we have to be clear about terminology for the case where there is no long transient for the deterministic skeleton of a model, yet the addition of stochasticity produces long term dynamics that are different than the equilibrium dynamics of the underlying deterministic model. As a way to outline the framework of the current paper, we summarize the current state of synthesis in Figure 1. The more systematic approach suggested by this figure first requires attention to the definition of transients and the kinds of stochasticity we consider, followed by different ways in which transients arise and the effects of stochasticity in different cases.

## 2 Simple examples of transients

Before starting with a systematic exploration of the influence of stochasticity on transients with an emphasis on long transients, simpler systems can provide background. Starting with the simplest case of linear deterministic systems, and then adding stochasticity, will demonstrate first the ecological importance of the phenomena, and provide insights into the role of stochasticity.

Age structured systems provide some of the simplest examples of transient dynamics, which are present even in linear systems. The dynamics of a population of salmon provide a straightforward illustration [38]. Individuals of most salmon species typically reproduce once and then die. In addition, in many populations, almost all individuals reproduce at the same age. We can denote the number of females of age i at time *t* by *n_i_*(*t*) in a discrete time description. We assume that the survival from age 1 to age 2 is given by *s*_1_ and similarly by *s*_2_ for age 2 to 3. Finally, denote the fecundity of 2-year-olds by *m*_2_ with m_2_ > 1 and the fecundity of 1- and 3-year-olds by *ϵ*_1_ and *ϵ*_3_ respectively, where *ϵ_i_* ≪ 1. Thus the dynamics of the females would be given by the following Leslie matrix model, if almost all individuals reproduce at age 2:

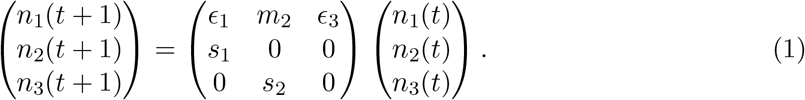

It is easy to see that if *ϵ*_1_ = *ϵ*_3_ = 0, this matrix would have two dominant eigenvalues of the same magnitude. If instead these fecundities are small and positive, then these two eigenvalues would have nearly the same magnitude. In this case, if in a given year almost all individuals were of age 2 and very few were of age 1, then for many years the dominant age class would alternate between 1 and 2. The addition of stochasticity could greatly reduce the time the system would need to approach stable age distribution (i.e. where the ratio of individuals in different age classes would be constant from year to year).

A second example of a linear ecological transient is given by a simple predator-prey system with an equilibrium that is a stable focus, but with complex eigenvalues with very small, negative real parts. In this case, the deterministic system would have oscillations whose magnitude would decay very slowly, while the presence of environmental stochasticity could extend the time to reach equilibrium [39] by interrupting the decay in cycle magnitudes.

What is interesting about these two simple examples is the contrasting effect of stochasticity. In the first one, the Leslie matrix model, clearly stochasticity would shorten the transient by accelerating the approach to the stable age distribution. In contrast, for the predator prey models, as originally demonstrated [40, 39] using Fourier analysis, a stochastic system can continue to exhibit cyclic behavior indefinitely and thus stochasticity greatly extends the lifetime of the transient, even making it effectively infinite.

If even linear systems can exhibit interesting and contrasting effects of stochasticity on transients, nonlinear systems, which can exhibit longer and more varied kinds of transients [8], will provide a much richer set of phenomena that will be key for ecological understanding. But, before presenting a systematic exploration of long nonlinear transients, it is important to highlight a particular class of stochastic transients which are prominent in ecology and more broadly.

Increasing attention is being paid to tipping points, and early warning signs for tipping have been developed based on the concept of critical slowing down [41, 42]. Critical slowing down describes the time a system takes to respond to a perturbation. Without stochasticity to perturb a system, there would be no opportunity to observe critical slowing down, and thus no possibility of early warning signs. Thus, the concept of critical slowing down has at its core ideas about both transients and stochasticity.

Transients also enter into another important aspect of early warnings. An emphasis of early warning sign work has been the detection of parameter changes that push the system through a saddle node bifurcation. At such a bifurcation, the system’s stable equilibrium is replaced by a ghost attractor which can lead to a long transient [7]. A natural question is how stochasticity affects the length (in time) of a transient resulting from a ghost attractor.

These simple but illustrative examples provide an important starting point for a discussion of transients in stochastic systems. But transients arise in many other ways, and given the ubiquity of stochasticity, a more thorough and systematic investigation is called for. Clearly, first steps are an unambiguous definition of transients, and a consideration of how stochasticity enters into ecological systems.

## 3 Definitions of long transients

There are two different ways to define transients in mathematical models (including those with noise) as well as in empirical systems. In this study we are emphasizing long transients due to their crucial role in ecological applications including sudden regime shifts.

Consider first the scenario where the system is functioning in a certain dynamical regime in which its major characteristics remain unchanged for a long time (for stochastic systems we operate with average characteristics). Here by ‘long time’ we understand the situation where the duration of the regime is much longer that its internal characteristic time (e.g. the period of oscillations). In an ecological context, this signifies that the duration of the regime is much longer than the generation time of species involved. To the external observer exploring the system based on time series, such a system would appear to be stable. Now suppose that at some moment in time, but without any changes to the properties governing the dynamics, the system demonstrates a rapid transition (as compared to the duration of the regime) to another regime which, in turn, conserves its new characteristics unchanged for a long time again. In this case, we call the preceding dynamical regime a long transient. Note that the post-transitional regime can be transient as well and a new transition may occur later on. In fact, the above mentioned scenario of transient behaviour describes a shift between regimes. It is also important to emphasize that according to the considered scenario the transition between regimes occurs without external forcing of the system, i.e. without changing model parameters in the course of time. Obviously, however, the presence of the long transient depends on the initial conditions for the system, so a regime shift due to long transients may be originally triggered by some initial disturbance of either the parameters or state of the system.

The other long transient scenario involves the situation where the system itself is in slow transition to a stable or quasi-stable state. We assume that the pattern of dynamics evolves very slowly with time as compared to the characteristic time of the current system. For example, this can be the case of damped oscillations with a very long relaxation time where both the amplitude and the period change only slightly. An important practical case is where the transition of the system to the final attractor actually requires an arbitrarily large time [31]. This can happen in the presence of large noise since there will always be perturbations kicking the system away from the eventual asymptotic state. In this case, the resultant pattern of dynamics will be an infinite sequence of transient regimes. We note that calling this behavior a transient is, perhaps, an arbitrary decision, as the combination of stochasticity plus the deterministic skeleton produces behavior that persists indefinitely. We believe that this is the more useful choice because it encompasses the role that stochasticity plays in altering dynamics and also note that this provides consistency in our definition.

Earlier [9] a key property of long transients was described: there is a scaling law describing the duration of transients while a particular model parameter is varied. The length of a transient regime (in stochastic systems the length should be understood as the mean length) can be made as large as possible when a certain bifurcation parameter (including the magnitude of noise) approaches a critical value. This mathematically quantifies the common sense notion of ‘long’ transient (i.e. how long is long). From the ecological point of view, the duration of a transient is always limited by natural constraints and we usually assume the average length of transients to be larger than several characteristic generation times [8]. The existence of a scaling law allows us to classify transients into different types [9].

## 4 Types of noise

In this contribution we focus primarily on extrinsic temporal noise, that is, those sources of stochasticity that arise because of relationships and quantities external to the modelled system. The classic example is, of course, detrended environmental variation. Coulson et al. [43] describe active and passive stochasticity, where active noise interacts with deterministic nonlinearity to produce dynamics that cannot result from either factor independently [44], and passive noise influences the transients among different deterministic states. The impact of environmental noise that affects the modelled system will depend on the modelled time frame of interest, the influence of the particular factor, the time scales of variation in the noise relative to the time scales of response, and the characteristics of the noise itself.

There are many properties to consider such as the whether the stochasticity is continuous, a single perturbation or seasonal, whether it has larger or smaller magnitude, and whether it has frequencies in a similar range as the intrinsic dynamics. Understanding the structure of noise is critical for understanding its potential impact. For example, continuous long term trended variation such as climate change, short term uncorrelated variation, and directed impacts through management might all be expected to have different effects on the same system.

The most familiar description of environmental stochasticity is as random draws, independent in time, from a Guassian distribution with small variance. In this white noise process, deviations from the mean at one timestep are unrelated to the size and magnitude of deviations at another timestep. That is, white noise is uncorrelated in time. Or put another way, all frequency components of the signal have the same expected value. The fact that the variance of noise is small has little bearing on its dynamical impact. To take a trivial case, even small variations close to a critical value in a bifurcation parameter can have large impact on the dynamics of a system. Environmental stochasticity of relatively small variance can also create oscillations through resonance effects [39, 45].

Of course most environmental signals such as temperature, rainfall and river flow rates have large variance and are autocorrelated in time even after detrending (e.g., [46, 47]). The strength of this autocorrelation depends on the signal itself (e.g., air temperature vs. sea surface temperature), the geographic location (e.g., continental air temperatures vs maritime air temperatures), and the time period. In particular climate change is altering the autocorrelation of related environmental signals [48, 49]. The autocorrelation of deviations can be large, and, in these cases, can cause clustering of extreme events [50]. Therefore, when we model the impact of environmental stochasticity as a white noise process, we may err in our estimates of the probability of long transient behaviour, as these signals can push a system away from (or towards) an attractor by virtue of the autocorrelated variation.

Of course larger amplitude noise, for example seasonal forcing [51, 52], can have large impacts, while episodic large amplitude noise (flow-kick) can move and maintain a system far from any attractor in the deterministic scaffold [53].

## 5 Interactions between stochasticity and transients

The effects of noise on transients are numerous and diverse (Fig. 1). Noise can make the lifespan of the transient considerably shorter and/or decrease the range of the initial conditions that result in the long transient dynamics, or remove the long transient altogether. Alternatively, noise can make the transient’s lifespan longer. With noise, the emergence of long transient dynamics becomes a probabilistic event rather than a deterministic one. Noise can turn a deterministic long transient into stable, persistent dynamics [40, 39]. Moreover, noise can create long transients via mechanisms that do not exist in a deterministic case [54, 55, 56].

The outcome of the interaction between noise and a long transient depends both on the properties of noise and on the mechanism behind the (deterministic) long transient. For the transient created by a crawlby [8] (i.e. caused by the closeness of the system to a saddle point), it is readily seen that uncorrelated noise makes the lifespan of the transient shorter (but does not remove it unless the noise is large), as the random movement of the system in the phase space pushes it, on average, away from the equilibrium. Consequently, the system does not necessarily follow the phase flow along the stable manifold that otherwise would bring it into the close vicinity of the saddle (cf. Fig. 2 in [9]). Interestingly, in the presence of noise, long transient dynamics can also emerge, with a certain probability, for a set of initial conditions that would not otherwise lead to a long transient, as the random movement of the system in the phase space can occasionally bring the system into close vicinity of the saddle.

**Figure 2:**
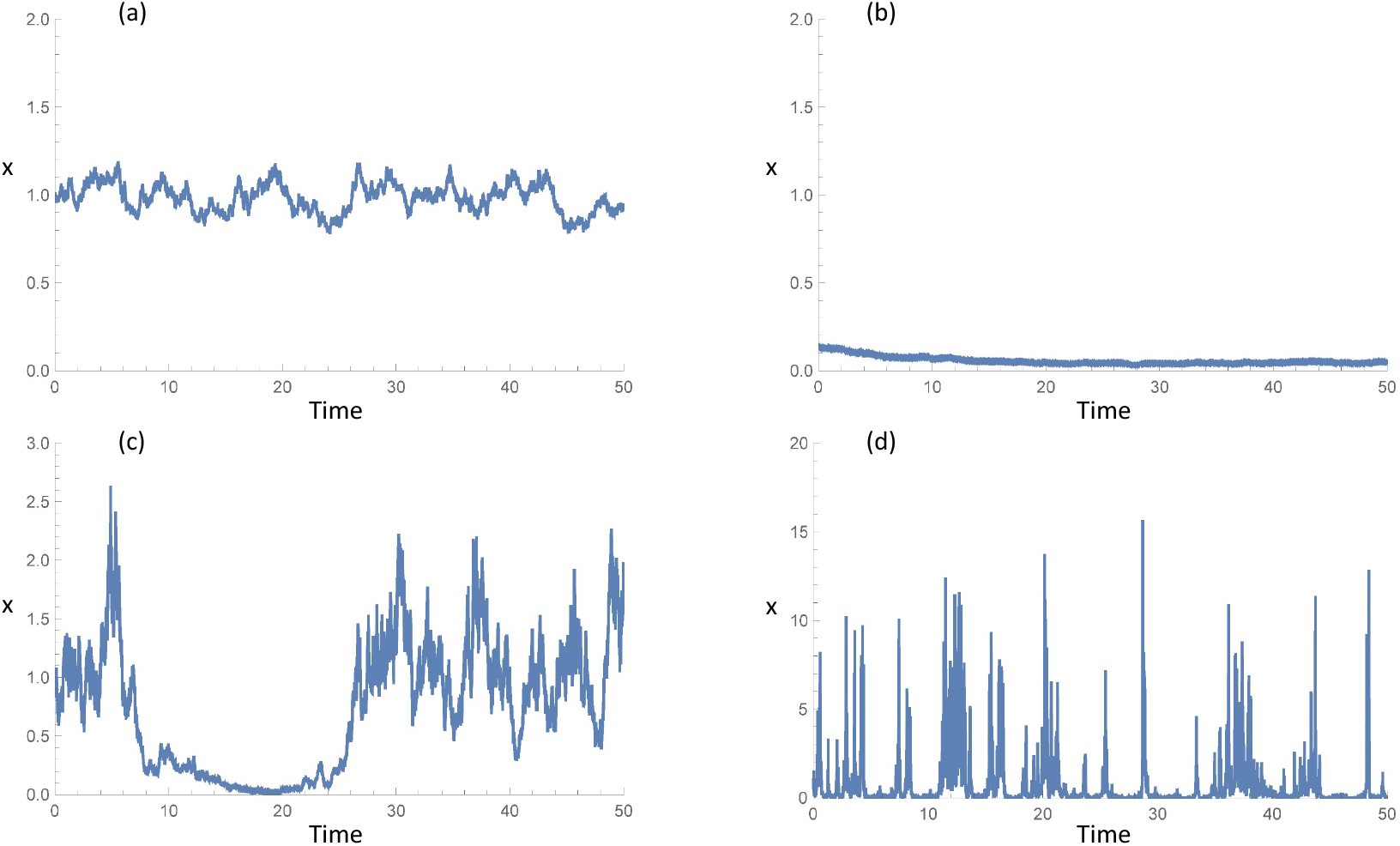
Behavior of a bistable ecological model with different noise levels. In each case the model with the deterministic skeleton *dx/dt* = *x*(*x* – 0.3)(1 – *x*) + 0.01, describing the dynamics of a population with scaled population size *x* describing Allee dynamics, is simulated for different noise levels. The deterministic skeleton is a bistable system where the last term represents a small steady immigration to prevent extinction. The noise term is of the form *γxw* where *w* is a Wiener process (white noise) with mean 0 and variance 1 and the equation is integrated as the equivalent Stratanovich stochastic differential equation using an Euler method and step size of 0.001. Note scale differences. (a) Small noise level, *γ* = 0.1, starting near the larger equilibrium; system stays near the equilibrium; (b) small noise level, *γ* = 0.1, starting near the lower equilibrium; system stays near the equilibrium; (c) intermediate noise level, γ = 0.6, showing noise induced transients; and (d) large noise level, *γ* = 5, where the system is noise dominated and does not exhibit transients.

The effect of correlated, directed noise can also make a transient much longer, by keeping the system in the vicinity of a saddle or ghost attractor. In particular, this is readily seen in a flow-kick system [57, 53] where the kicks (directed, quasi-periodical, time-discrete random perturbations of the state variable) control the movement of the system over the phase space, with the capacity of keeping it close to a specific location in that space (e.g. a saddle or a ghost attractor).

Perhaps the simplest and best known example where noise can change the system properties qualitatively is the bistable system. We mention here that bistable systems are highly ecologically relevant; in particular, they are used as the paradigm of a regime shift [58] resulting from slow parameter change. Without noise or even with extremely small noise, the system remains in the vicinity of one of the steady states indefinitely long (see Fig. 2a,b). However, slightly larger uncorrelated noise can push the system out of the attraction basin of the current state, so that it fast converges to the alternative state: a purely noise-induced regime shift occurs. The dependence of the state variable (e.g. the population size) on time takes the form of alternating periods with a quasi-stationary value (Fig. 2c). The time spent by the system in the vicinity of the given state is inversely proportionate to the noise intensity and, hence, can be very long. Therefore, small noise creates long transient dynamics. We mention here that there are empirical examples of stochastic switching with long transients in ecological systems [59, 60, 61] as well as in epidemiology [62]. Interestingly, noise of larger intensity can destroy the long transient as the system diffuses across the whole span of the phase space between the two states (Fig. 2d). Therefore, the dependence of the lifetime of the transient dynamics on the strength of noise is non-monotonous. This non-monotonicity is a generic property of population dynamics with stochasticity and is seen in a variety of systems and models (e.g. see [63, 64, 65]). Below, we see a similar phenomenon emerging in high dimensional ecological systems.

Another example of a situation where noise can create long transients is the dynamics of excitable systems [55]. A relevant ecological system that exhibits excitable dynamics is a prey-predator system with Holling type III predation [66]. In a certain parameter range (e.g. where the linear predator nullcline is to the left of the trough of the prey nullcline, see Fig. 3a), the coexistence state is globally stable, but there is a threshold separating different types of approach to it. For initial conditions on one side of the threshold, the system approaches the steady state directly. For initial conditions on the other, excitable side of the threshold, the system takes an excursion around state space to large abundance of prey and then predator before returning to settle at the steady state. In the deterministic case, once the system has returned to the steady state, it stays there indefinitely. However, the excitability threshold runs close to the steady state, so noise can push the system over the threshold, triggering another large excursion around the phase plane before finally returning to the vicinity of the coexistence state where it can remain for a long time until noise pushes it out again (see Figs. 3b,e). Altogether, the state variable exhibits small-amplitude, random oscillations around the steady state value intermittent with occasional large-amplitude cycles. An increase in the noise level makes the large-amplitude cycles more frequent, see Figs. 3c,f. The periods of small-amplitude oscillations are long transients. This dynamic can be viewed as a noise-induced mixed-mode oscillation [67, 68].

**Figure 3:**
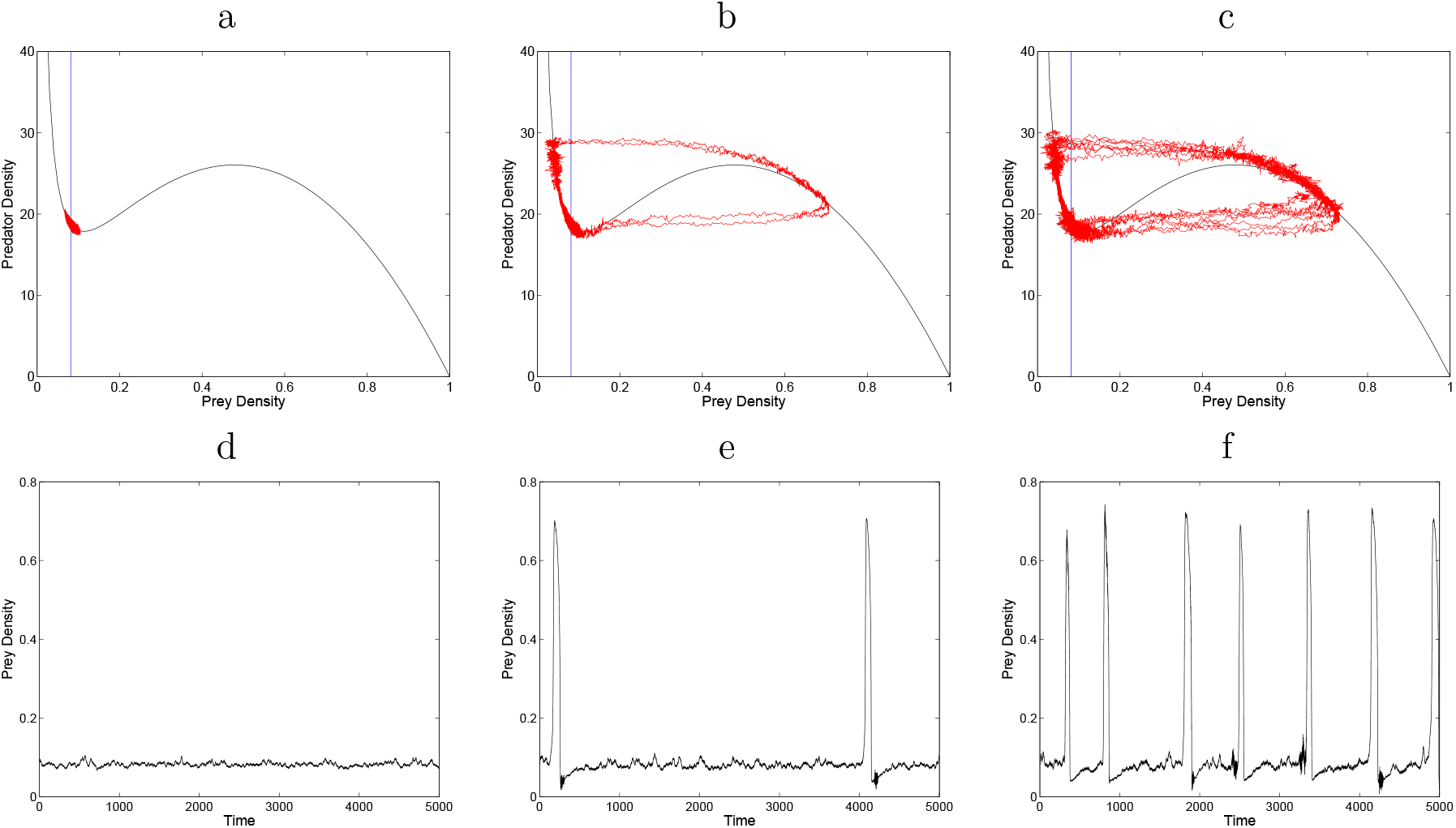
Excitable system under the effect of linear additive noise. (a,b,c) The phase plane of prey-predator system with Holling Type III predation: the irregular red curves show the system trajectories for different level of noise (increasing from a to b to c), the vertical line is the predator nullcline, the S-shaped curve is the prey nullcline. (d,e,f) The corresponding dependence of the prey density on time.

Another highly relevant example of transient dynamics facilitated by noise is noise-induced synchronization [56, 69]: population oscillations at different locations in space (e.g. in different patches of a fragmented habitat) that would occur asynchronously in the absence of noise can become synchronized under the effect of noise. In ecology, this phenomenon is often referred to as the Moran effect and it is believed to be responsible for masting [70, 71]. However, full synchronization only happens when the controlling parameter (e.g. the strength of the noise) exceeds a certain critical value. In the subcritical parameter range, intermittent synchronization occurs, so that the periods of synchronized and asynchronized dynamics alternate [72, 56]. In this case, the intervals of synchronized dynamics can be regarded as transients. When the controlling parameter approaches its critical value, their lifetime becomes very long; the average transient time follows the power law [72].

Noise can also turn a transient regime into permanent, sustainable dynamics. As a simple example, let us consider damped population oscillations. In models, such oscillations are frequently observed around a stable focus. Their characteristic life time is *τ* ~ 1/ |Reλ_0_| where λ_0_ is the eigenvalue with the largest negative real part. Correspondingly, for |Reλ_0_| ≪ 1, they last for very long and hence can be regarded as long transient dynamics. The effect of noise can be to turn these long-term damped oscillations into a sustained oscillations [40, 39, 73] through a mechanism known as stochastic resonance [74]. Such quasi-cycles have been reported in several empirical systems including the dynamics of populations of Dungeness crab [31] and bluefin tuna [75].

As we noted above, tipping points with simple saddle node bifurcations of equilibria are a core example of the interaction between stochasticity and transients. This interaction becomes even more important with a deterministic system with a chaotic attractor that experiences a crisis [76] when its controlling parameter *p* passes the critical value *p_c_*. Before the bifurcation point, i.e. for *p* < *p_c_*, there is a chaotic attractor so that chaos is self-sustained; at *p* = *p_c_* the chaotic attractor turns into a chaotic saddle so that for *p* > *p_c_* chaotic dynamics are transient. In the presence of chaos, there is a non-zero probability that the system will leave the basin of the attractor; hence the sustainable chaotic dynamics become transient rather than sustained. The classic three species food chain [77] is an important ecological example that has this kind of bifurcation. The presence of noise further complicates the situation and can lead to very long transients, supertransients, even for those parameter values where the deterministic system would have a stable chaotic attractor. As previously reviewed [9], in the region where there is deterministic stability, after some time the corresponding stochastic system can cross the basin boundary and leave the basin of attraction for the chaotic attractor and end up in the basin of attraction for a different attractor. This situation is thus a case where stochasticity leads to a transient. A rigorous mathematical analysis of this case is possible [78, 79].

## 6 High-dimensional ecological systems

There is one more important class of systems where stochasticity plays an important role in transients, namely high-dimensional systems. The study of high-dimensional stochastic systems is important because ecological systems are typically high-dimensional, nonlinear and complex. The high dimension can arise either from interactions among many species, or simply from including explicit space in the description. The study of such systems is inherently complex and in its early stages, so rather than give a comprehensive guide, instead we focus on an example that is particular illustrative. A representative class of systems is mutualistic networks [80, 81, 82, 83, 84, 85, 86, 87, 88, 89, 90], e.g., a bipartite network of pollinator and plant species. Because the number of species involved in the mutualistic interactions can be large, the system is high dimensional.

Consider a complex mutualistic network subject to environmental or demographic noise, or both. The setting thus naturally has high dimensionality and stochasticity. Can transients arise and are they typical? The answer is affirmative. One scenario is tipping point dynamics [91, 41, 42, 92, 93, 15, 94, 95, 96, 97, 98, 99, 83, 100, 101]. In particular, in a mutualistic network, the deterministic behavior is dominated by the dynamics about a tipping-point transition [87, 88]. For example, environmental deterioration will result in massive species extinction, which can occur suddenly in mutualistic systems as a relevant parameter (e.g., the species decay rate) increases through a critical point – a tipping point. Under noise, even when the parameter value has not reached the tipping point, a total system collapse can occur. This is the phenomenon of noise-induced collapse which, dynamically, is nothing but a transition from one steady state to another: from a healthy, high-abundance state to an extinction state. The collapse, of course, does not occur instantaneously: it takes time for the transition to complete, and during this time what we see is a transient. Likewise, when the system is effectively extinct with near zero species abundances, noise can trigger a recovery of the species abundances. In this case, the transition occurs in the opposite direction: from a low abundance steady state to a high abundance one, which is the recently discussed phenomenon of noise-induced recovery [89] accomplished through a transient.

A dynamical picture of the phenomenon of noise-induced collapse and recovery is illustrated in Fig. 4. In the deterministic case, species collapse and recovery are the result of saddle-node bifurcations. Let *κ* be the normalized species decay rate (the bifurcation parameter). Environmental deterioration is manifested as an increase in the value of *κ*. As *κ* increases through a critical point, denoted as *κ_c_*(0), a reverse saddle-node bifurcation occurs, giving rise to a tipping point transition. Now consider the case where noise of amplitude *ε* is present. The phenomenon of noise-induced collapse corresponds to an earlier tipping point transition, now occurring at the critical point *κ_c_*(*ε*), where *κ_c_*(*ε*) < *κ_c_*(0). Likewise, without noise, species recovery occurs through a forward saddle-node bifurcation at *κ_r_* (0), but noise can induce species recovery at a critical point *κ_r_*(*ε*), where *κ_r_*(*ε*) > *κ_r_*(0). For *κ_r_*(0) < *κ* < *κ_c_*(0), the deterministic system has three equilibria: two stable equilibria and an unstable equilibrium in between. The two stable equilibria are two attractors with their own basins of attraction, while the stable manifold of the unstable equilibrium is the basin boundary [102, 12]. Dynamically, the two transition phenomena are the result of noise driving the system across the basin boundary. Transients arise because of the competition between the attractive dynamics in the neighborhoods of the stable equilibria as controlled by the eigenvalues of the Jacobian matrix with negative real part, and stochastic hopping that brings the system out of the attractor [103, 104, 105]. The transient dynamics underlying noise-induced collapse and recovery are the result of stochastic forcing that drives the system from one stable steady state to another. For an ensemble of trajectories from random initial conditions, the transient time required for the transition is typically exponentially distributed [106, 107] and the average transient lifetime *τ* depends on the noise amplitude *ε*. For stronger noise, the transition occurs more quickly, so we expect *τ* to decrease with *ε*. A recent study of four real world mutualistic networks [108] demonstrated the phenomena of noise-induced collapse and recovery, and confirmed the occurrence of transients. Computation with a two-dimensional reduced model [87] of the empirical networks revealed [108] the following algebraic scaling law between *τ* and *ε*:

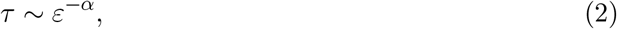

where *α* > 0 is the algebraic scaling exponent.

**Figure 4:**
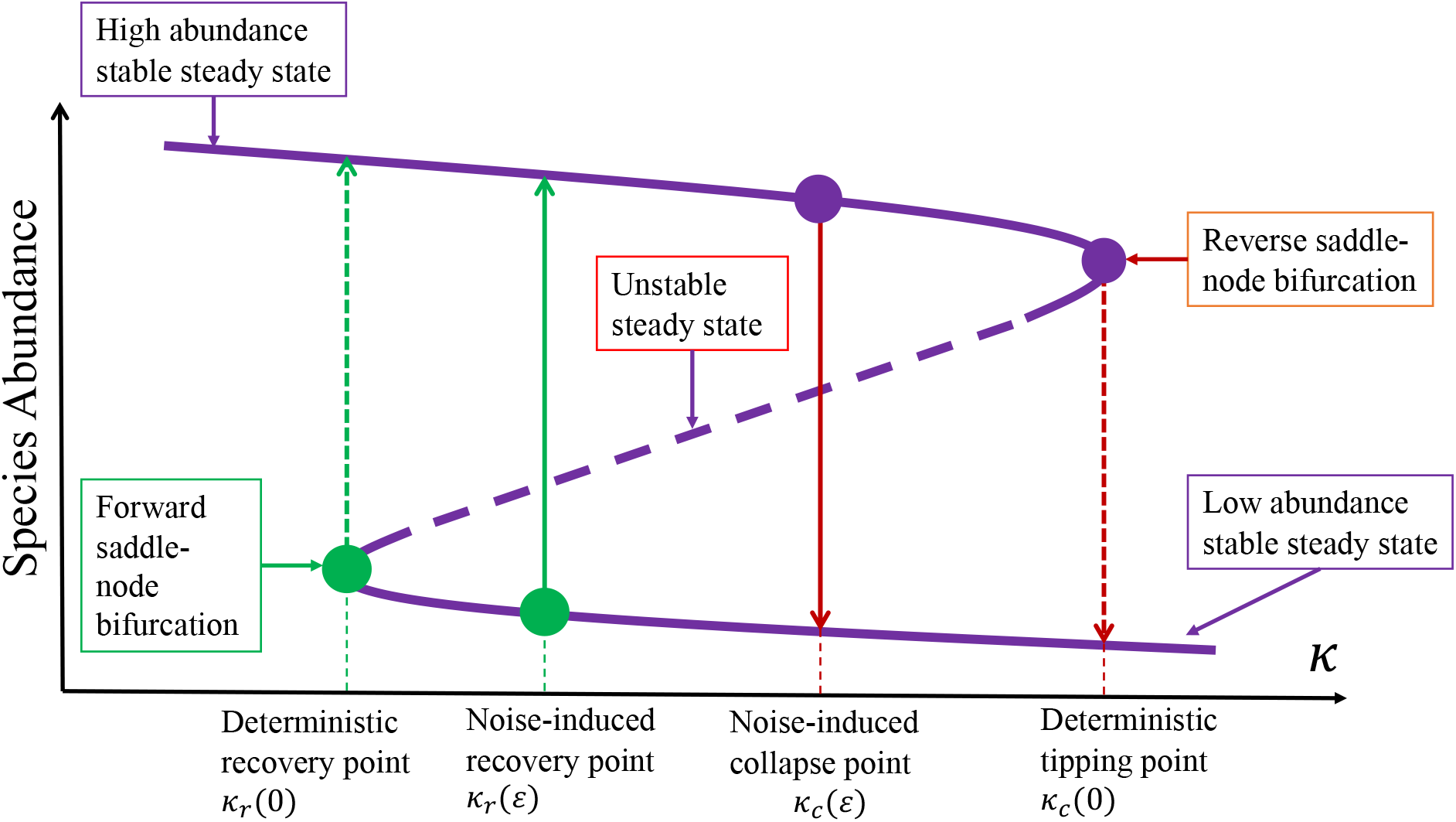
Dynamical mechanism of noise-induced collapse and recovery in mutualistic networks. The bifurcation parameter is the species decay rate *κ*. The system has two stable steady states, with high and low abundances, respectively. There is an unstable steady state in between the two stable states. The three equilibrium points are dynamically connected through two saddle-node bifurcations: one corresponding to the tipping point (the reverse one) at *κ_c_*(0), and another leading to species recovery (the forward one) at *κ_r_*(0). In the deterministic case, as *κ* increases through *κ_c_*(0), the system collapses. Under noise of amplitude ε, the collapse can occur earlier at *κ_c_*(*ε*) – the phenomenon of noise-induced collapse. Likewise, as κ decreases, noise can induce species recovery at *κ_r_* (*ε*).

The common dynamical feature of transition from one stable steady state to another between noise-induced collapse and recovery notwithstanding, the specific nature of the noise does play an important role. In particular, environmental noise is independent of the dynamical variables of the system and is thus simply additive, but demographic noise depends on the species abundances. Before reaching the tipping point where the system is in the high-abundance steady state, demographic noise is strong and is the dominant stochastic force to induce a system collapse. In contrast, if the system is in the low-abundance steady state, demographic noise is weak. In this case, the environmental noise becomes the major stochastic driving force for system recovery. The specific roles played by demographic and environmental noises have implications to devising strategies to manage high-dimensional ecological systems. For example, because of the detrimental role of demographic noise in causing an ecosystem to collapse, it is imperative to devise methods to reduce the level of demographic noise to keep the system in the healthy state. Conversely, when the system is already close to extinction, a suitable amount of environmental noise may facilitate recovery [89].

## 7 Flow-kick dynamics and single realizations

So far, we have focused on systems where the stochastic influence is due to continual noise. But in real ecological systems there may instead be large disturbances at regular or irregular intervals. These exogenous disturbances may be stochastic or, as in some management settings, they may be tightly controlled. In analyses of transients in a deterministic setting [8, 9], the focus is often on response to a single perturbation of a system away from its asymptotic state. These ideas provide the background behind the approach we use to deal with cases of repeated large disturbances.

Consider a population that – in the absence of noise – has an attracting state. If the system is subject to recurring disturbance by disease, weather extremes, management, etc., it may never even get close to the asymptotic dynamics, but instead will stabilize in a region of state space where the short-term transient dynamics balance the disturbance.

A familiar example is given by fishery management: an undisturbed fish stock might grow to carrying capacity but, when subject to repeated harvesting, does not recover to full carrying capacity between harvests. Indeed, a management strategy typically maintains stock population significantly below carrying capacity to ensure a high recruitment rate and corresponding yield. Alternatively, in the context of an invasive species, the disturbance pattern might represent a culling strategy. In this case a management strategy of regular removal may be designed to keep the invasive below a threshold way below carrying capacity.

One approach to exploring this phenomenon mathematically is to combine the growth dynamics of the ecosystem with the disturbance dynamic to define a new system whose asymptotic dynamics represent a balance between growth and disturbance. For example, given a continuous population model 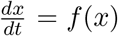 that is repeatedly disturbed by a discrete kick *κ* to the state variable, the associated flow-kick system (a special case of impulsive differential equations), is defined by the discrete system

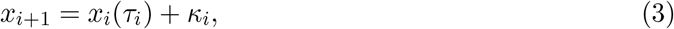

for additive disturbance, or

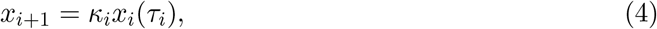

for multiplicative disturbance. Here *x*(*t*) is the solution to the undisturbed system with initial condition x, and the ith kick *κ_i_* occurs at time *τ_i_* after the previous kick *κ*_*i*−1_. In the fishery context, *f*(*x*) represents the recruitment function, and the disturbance pattern *κ_i_,τ_i_* represents a harvesting strategy. This framework is used in [57] to quantify resilience of ecosystems to regular recurrent disturbances, and in [53] to study the resilience of socially valued properties of natural systems to recurrent disturbance.

This kind of flow-kick system illustrates the essential role played by transient dynamics in the presence of disturbance to yield different qualitative dynamics, in which kicks can move the asymptotic state of the system in any arbitrary way [53]. More formally, given any point *x*\* in n-dimensional state space, and any disturbance time *τ*, there is a kick *κ* so that *x*\* is an equilibrium of the flow-kick system (3) with *κ_i_* = *κ* and *τ_i_* = *τ* for all *i*. In other words, using a perfectly regular disturbance pattern of fixed kicks at fixed time intervals, one can stabilize the disturbed system *anywhere* in state space, regardless of the location of attracting sets or basins of attraction of the underlying growth dynamics 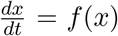. This idea can be a very powerful conceptual tool in the management of ecosystems [109].

## 8 Conclusions

An overarching challenge in understanding the dynamics of ecological systems is to provide insights on ecologically-realistic time scales in the presence of both environmental variability and stochasticity driven by small population sizes. In this setting, the asymptotic behavior of deterministic systems is not relevant, and instead a focus on transient dynamics is required. Examples of transient behavior have been observed in a variety of ecological systems as previously summarized [8], but a more systematic approach is important for understanding the role of stochasticity in ecological transients, especially in cases where detailed information about the system may be limited. The importance of transients in stochastic systems shows up in a variety of ecological contexts [10] and our contribution here emphasizes both the importance of this phenomenon and the idea that a careful mathematical treatment can find order in what may seem like a series of idiosyncratic examples. This is an area where despite the advances we have covered here, much more work is needed.

The work summarized here can also be thought of as an extension to stochastic variation of the insights that come from studying seasonal dynamics [52]. Analyses of seasonal systems have emphasized ecological implications, such as long-term variation of densities and species succession of plankton communities of temperate lakes across the warm season [110], where the random starting conditions play an important role. The existence of transients with time scales much larger than the time of a single season can guarantee the coexistence of many plankton species within a short time period – which would be impossible for a longer period in a constant environment – since the ecosystem is ‘re-set’ each year in a random fashion [111]. In other words, this type of ecological transient seems to be a robust phenomenon; however, mechanisms of observed long transients in many such systems are still unclear due to the high complexity of communities containing dozens of interacting species, the existence of several time scales, and stochastic aspects. Further work based on the ideas we have developed here will shed light on the ecological implications of stochastic transients [10].

Here, we have emphasized using modeled stochastic dynamics as a way to understand and predict real-world dynamics. We might also consider the inverse problem, in which we could think of stochasticity as obscuring the signal of a processes of interest. As a result, it is tempting to feel that ecological insights would be improved if we could study ecological systems in isolation from stochastic noise. However, if we could observe ecological dynamics in the absence of noise, we would see one behavior – the steady state. A system at or near its equilibrium would simply sit at equilibrium. If our observations began with the system out of equilibrium, we could see a short transient or part of a long transient eventually approaching the equilibrium. In the presence of noise, however, we have the opportunity to see all of these things within a reasonable observation window, as perturbations push the system from one domain to another [112].

One key example that we have highlighted in this work is unexpected shifts between alternative stable states in ecological systems [41]. Much attention has been focused on single shifts, but many systems move more than once between different states. Important quantities, like the expected interval between shifts and the proportion of time the system is expected to be in each state, can be computed with knowledge of each stable state’s basin of attraction [113] and the characteristics of the noise. A system that mostly sits at or very near one equilibrium gives us virtually no information about these basins. We may not even know whether other stable states exist in such a system. In contrast, a system that experiences enough stochasticity could shift many times [65].

In conclusion, here we argue that an investigation of noisy nonlinear systems reveals a much richer view of the underlying *deterministic* structure of ecological systems than does focusing exclusively on unperturbed systems. When we observe a system at equilibrium, we can only infer that the equilibrium exists, not what causes it. When we observe how different parts of the system – such as the population densities of different interacting species – change in response to being in different states or configurations, we gain valuable information about the nonlinearities and feedbacks that are present. Extracting these insights, however, requires a good understanding of how the types of stochasticity present interact with these nonlinearities and feedbacks.

Finally, we emphasize that although we have focused on ecological issues and models here, these themes arise in other areas as well. In particular, interactions between stochasticity and transients are clearly important in neuroscience [114, 115], as well as other areas of biology, engineering [116], physics [117] and climate.

## Acknowledgements

This work was conducted as part of the Long Transients and Ecological Forecasting Working Group at the National Institute for Mathematical and Biological Synthesis, supported by the National Science Foundation through NSF Award DBI-1300426, with additional support from The University of Tennessee, Knoxville, and NSF Award no. CCS-1521672. The long term support of NIMBioS directors Lou Gross and Sergey Gavrilets is greatly appreciated. The work of AH was also supported by NSF Grant EF-2025235, and of MLZ by NSF Grant CCF1522054.

